## Supplementary materials for "Real-time spectroscopic photoacoustic/ultrasound (PAUS) scanning with simultaneous fluence compensation and motion correction for quantitative molecular imaging"

to the paper

**TABLE OF CONTENTS**

**Supplementary Note 1**. Diode-pumped, wavelength-tunable laser.

**Supplementary Note 2**. PAUS transmission/detection head.

**Supplementary Note 3**. Flow chart for spectroscopic PA imaging.

**Supplementary Note 4.** Σλ-PA and component-weighted imaging.

**Supplementary Note 5.** Agent properties.

**Supplementary Note 6.** Light-diffusion based laser fluence estimation in the fast-sweep geometry

**Supplementary Note 7.** Wavelength-dependent fluence estimation in 1%, 2%, and 4% Intralipid.

**Supplementary Note 8.** *ex vivo* fluence compensation results in chicken breast

**Supplementary Note 9.** Motion estimation using PatchMatch

**Supplementary Movie 1**. Real-time *ex vivo* multispectral imaging for needle guidance with nanoparticle injection into chicken breast

**Supplementary Movie 2**. Real-time *in vivo mouse* multispectral imaging for needle guidance with nanoparticle injection.

**<u>Supplementary Note 1</u>. Diode-pumped, wavelength-tunable laser**

A unique, compact (see Specification sheet below), diode-pumped, high pulse repetition rate (from single shot to 1000 Hz), wavelength tunable (700-900 nm) diode-pumped laser (Laser-Export, Moscow, Russia) was designed and built especially for the fast-sweep PAUS system.

The laser can emit pulses of about 1 mJ energy at a 1000 Hz repetition rate, with wavelength switching times less than 1 ms for any arbitrary order of wavelengths (i.e., wavelength does not need to be sequentially stepped between bounds). Every laser pulse in a sequence can be at a different wavelength without sacrificing the kHz repetition rate. The number of pulses per wavelength in a sequence and energy of every pulse in percent from maximum can be pre-set. The pulse repetition rate can be set in the range 1 – 1000 Hz for both single- and multi-wavelength operation. Other laser parameters are listed in the specification sheet below.

### LASER SPECIFICATION

Wavelength tuning range .......... 700 - 900 nm
Mode of Operation .......... Q-switched
Average Pulse Energy at 1 kHz at 700 nm .......... > 500 μJ
Average Pulse Energy at 1 kHz at 900 nm .......... > 600 μJ
Max. Average Pulse Energy at 1 kHz in the range of 780 - 820 nm .......... 1000 μJ
Beam Profile .......... $TEM_{00}$
Beam Diameter ($1/e^2$, at output aperture) .......... 0.9 mm
Pulse duration at the wavelength of maximum energy .......... 8-13 ns
Pulse-to-Pulse Stability (StdDev/Mean, 1 kHz, at nominal pulse energy)
    at set wavelength regime .......... <8 %
    at varying wavelength regime (for pulses of the same wavelength) .......... <10 %
Polarization Linearity .......... > 100:1
Range of Pulse Repetition Rate:
    Ext. Triggering .......... 0 - 1 kHz
    Int. Triggering: through RS-232 .......... 0 - 1 kHz
    without PC .......... 1 ± 0.01 kHz
Trigger Pulse Fall Egde (or Q-switcher trigger pulse) to Laser Pulse
    Delay (the value within the range) .......... 300 ± 250 ns
    Jitter .......... < 8 ns
Dimensions (maximum linear size ):
    Laser Head .......... 250 mm
    Power Supply .......... 300 mm
Weight:
    Laser Head .......... <6 kg*
    Power Supply .......... < 7.5 kg*
Wavelength Switching Rate per Step .......... <1 ms
Wavelength Tuning Control .......... preset mode, PC mode
Remote Control of Laser Parameters:
Pumping ON/OFF, Ext./Int. Triggering, pulse repetition rate, pulse energy .......... via RS-232
DTR via RS-232 .......... 4800 bit/s
Operating Temp/Humidity Range .......... +15 to + 35 °C /up to 80% non-condensing
Operating Voltage .......... 24 ± 10% VDC
Laser Class .......... IV
Compliance .......... CE, RoHS

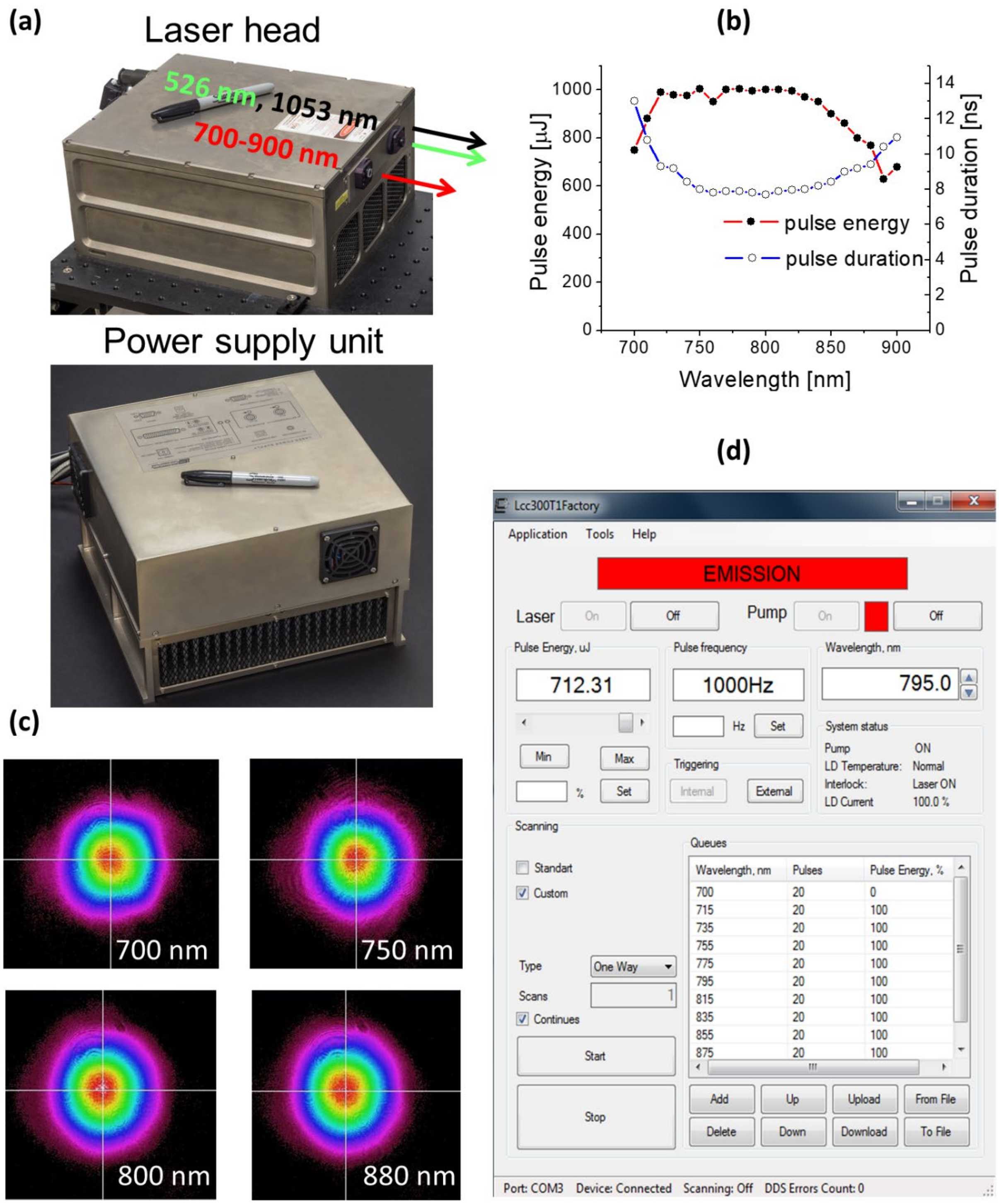


**Suppl. Fig. 1: Diode-pumped, kHz rate, wavelength-tunable laser. a,** photograph of the laser; **b**, wavelength dependence of laser emission energy and pulse duration; **c**, beam profiles; **d**, an example of a pulse sequence (implemented continuously with no delays for wavelength or regime switching).

**Supplementary Note 2. PAUS transmission/detection head**

The PA probe head combines a commercial 15-MHz, 128-element US linear-array probe with a pitch of 0.1 mm (Vermon, France) with 20 fibers spanning the elevational extend of the array as two fiber arrays (Supplementary Fig. 2a). Each fiber array (i.e., 10 fibers) has an interval of 1.5 mm, spanning 13.5 mm over the whole US lateral image range (i.e., 12.8 mm). Two fiber arrays are separated by a distance of 12 mm, and tilted symmetrically relative to the US image plane with an angle of 35°. Consequently, beams generated from each fiber array intersect on the same image plane at a depth of 8 mm around the transducer's elevational focus (Fig. 2b). The impulse and frequency responses of the whole system, including the probe to PA acquisition, were measured by receiving the PA signal at a single channel from a point-like absorber (i.e., human hair), as shown in Fig. 2d.

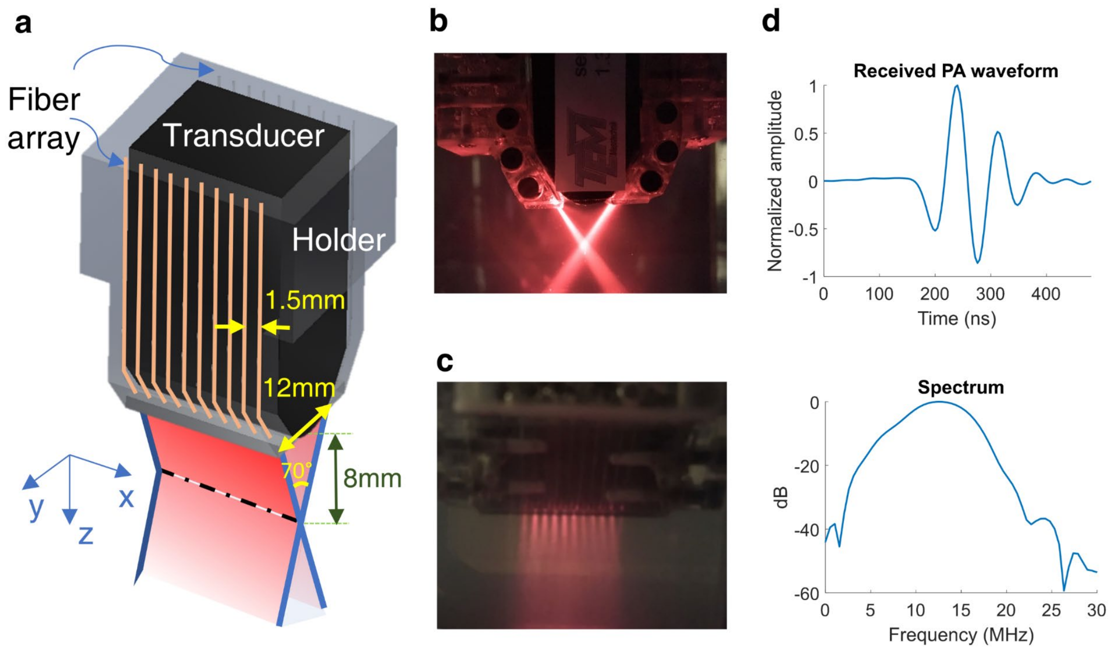


**Suppl. Fig. 2: Integrated PA probe with 20 fibers**. **a,** 3-D schematic view. **b,** photograph of the integrated probe viewed in the y-z plane. **c,** photograph viewed in the x-z plane. **d,** Measured PA waveform and frequency response from a point-like absorber (i.e., human hair).

**Supplementary Note 3**. **Flow chart for spectroscopic PAUS imaging**

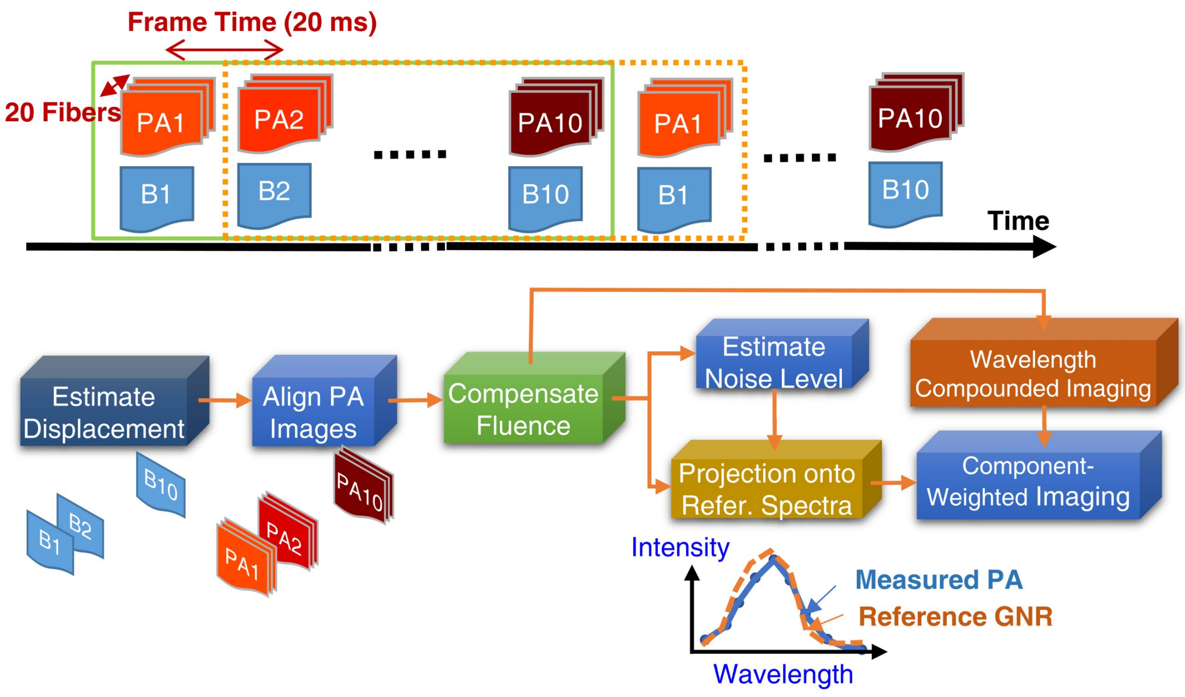


**Suppl. Fig. 3: Flow chart for spectroscopic PAUS imaging using**. Ten (10) different laser wavelengths form a spectroscopic scan cycle (see the green solid rectangle). Motion among all 10 B-modes is estimated. The estimated displacement is then applied to all PA sub-images. By identifying strong absorbers in a PA image, wavelength-dependent optical fluence is estimated using the diffusion approximation. The estimated fluence is then compensated over all PA sub-images over the 10 different wavelengths. A wavelength-compounded imaging (called $\Sigma\lambda$-PA) is then produced by coherent summation over all 10 wavelengths. Quantitative PA spectroscopy is performed pixel-by-pixel. To accurately estimate a component spectrum, the noise level is estimated by turning off the laser at 700 nm and then subtracted from the measured spectrum. The compensated, noise-subtracted spectrum is projected onto the reference spectrum (e.g., the absorption spectrum measured from a UV-VIS spectrophotometer). Finally, multiplying $\Sigma\lambda$-PA by the projected result produces component-weighted imaging. By shifting one-wavelength frame, another new spectroscopic image can be obtained simply using the latest 10 wavelengths at a 50 Hz frame rate, after appropriate motion and fluence compensation over this set of measurements (see the yellow dashed-line rectangle). Note that a persistence filter can be applied to produce smooth spectroscopic results at a reduced rate but the spectroscopic frame is updated at 50 Hz.

**Supplementary Note 4**. **Σλ-PA and component-weighted imaging**

Both wavelength-compounded PA (Σλ-PA) and component-weighted PA images are processed after performing motion correction and fluence compensation. As shown in Supplementary Fig. 4, Σλ-PA images are the coherent sum over all wavelength PA images (i.e., 9 wavelengths ranging from 715 to 875 nm used in this study). Component-weighted PA images are produced by projecting specific molecular constituent spectra onto the measured PA signal spectrum.

A projection is obtained by computing the correlation between the measured PA spectrum and the known spectrum of a specific molecular absorber. First, all 9 wavelength PA images are subtracted pixel-wise from noise estimates obtained by turning off the 700 nm laser emission. Next, for the $i$-th pixel, the normalized cross correlation (NCC) between the spectral estimate $x_i(\lambda)$ from $N_\lambda$ wavelengths and the reference spectrum $y(\lambda)$ (i.e., UV-VIS measurements) is computed:

$$C_{x_i,y} = \frac{\sum_{k=1}^{N_\lambda}[x_i(\lambda_k) - \bar{x_i}][y(\lambda_k) - \bar{y}]}{\sqrt{\sum_{k=1}^{N_\lambda}[x_i(\lambda_k) - \bar{x_i}]^2}\sqrt{\sum_{k=1}^{N_\lambda}[y(\lambda_k) - \bar{y}]^2}}, \tag{S1}$$

where $C_{x_i,y}$ is the projected value for the $i$-th pixel. $\bar{x_i}$ and $\bar{y}$ represent the mean value of the estimated and reference spectrum over all $N_\lambda$ wavelengths, respectively.

Using the NCC for a particular absorber, a weight is obtained at each pixel in the image for that absorber from a scaling function. In this study, a sigmoid was used to produce the weighting map for each absorber to minimize noise for low NCC and highlight regions of high NCC. For each NCC value *X*, the weighting result *Y* is given by

$$Y = \frac{1}{1 + e^{-a(X-b)}}$$
$$0 \le X, Y \le 1;\;\; 0 \le a \le \infty;\;\; 0 \le b \le 1\;, \tag{S2}$$

where *b* represents the NCC value corresponding to *Y* = 0.5 and *a* determines the slope of the function (Supplementary Fig. 4). This non-linear function can slightly distort quantitative information in pixels where multiple absorbers contribute to the signal, but it maintains quantitative accuracy in pixels dominated by a single absorber while suppressing noise in regions of low PA signal. In this study, *a* and *b* for GNR-weighted PA images are 50 and 0.82, respectively. *a* and *b* for needle-weighted PA images are 60 and 0.9, respectively. Finally, a component weighted PA image is produced for each molecular absorber by multiplying the appropriate weighting map with the Σλ-PA image on a pixel-by-pixel basis.

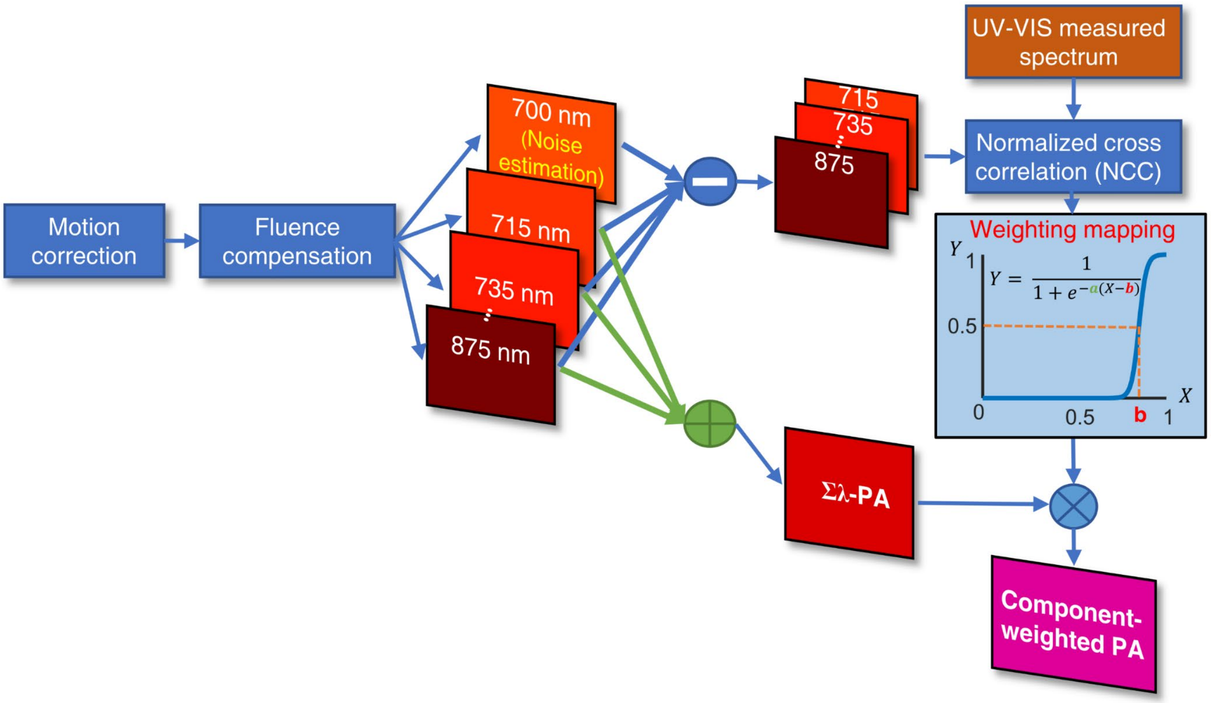


**Suppl. Fig. 4:** Signal processing flow chart for wavelength-compounded (Σλ-PA) and component-weighted PA imaging.

**<u>Supplementary Note 5</u>. Agent properties**

**Gold nanorods solution.** PEG-coated, 45 nm long, gold nanorods (GNRs) manufactured by NanoHybrids (Austin, TX, USA) were used. The properties are summarized in Supplementary Table 1. The measured UV-VIS absorption coefficient of the GNR solution is shown in Supplementary Fig. 5a. Given the size and aspect ratio of the GNR, the solution optical density is dominated by optical absorption.

**Supplementary Table 1.** Properties of gold nanorods used in this study

| | |
|---|---|
| **Longitudinal Peak** | **776 nm** |
| **Longitudinal Optical Density** | 51.0 |
| **Peak Width at 80%** | 68 nm |
| **Transverse Optical Density** | 12.4 nm |
| **Longitudinal/Transverse** | 4.1 |
| **Particle Concentration** | $3.9\times10^{13}$ per ml |
| **Mass Concentration (Au)** | 2.2 mg/ml |
| **Width** | 11.4 ± 1.1 nm |
| **Length** | 44.8 ± 3.4 nm |
| **Aspect ratio** | 3.9 |
| **Solvent** | De-ionized and ultrafiltrated water |
| **Particle Surface** | mPEG, 5kDa |
| **Zeta Potential** | -3.1 mV |
| **pH** | 7.1 |

**Black ink solution.** Black ink solution (#44011, Higgins, MA, USA) was used as a strong absorber injected into the tube without dilution. The measured extinction spectrum is shown in Supplementary Fig. 5b where 30 µl black ink was diluted by 1 ml de-ionized water.

**Prussian blue nanoparticles.** Prussian blue nanoparticles were synthesized using methods adapted from Zhu *et al.* [1] and Ren *et al.* [2] To prepare the Prussian blue nanoparticles, stock solutions of 250 ml of 1 mM iron (III) chloride (Sigma-Aldrich, product number: 157740, CAS: 7705-08-0), 250 ml of 1 mM potassium ferrocynide (Sigma-Aldrich, product number: P3289, CAS: 14459-95-1), and 500 ml 0.5 mM sodium citrate solution (Sigma-Aldrich, product number: W302600, CAS: 6132-04-3) were prepared in pure 0.2 µm filtered de-ionized water. All stock solutions were brought to 60°C while stirred using a magnetic bar while on a hotplate. The sodium citrate solution was divided equally between the iron (III) chloride solution and the potassium ferrocynide solution and stirred until fully mixed. Finally, the potassium ferrocynide with sodium citrate solution was added to the iron chloride with sodium citrate solution, gradually forming a slightly green tinted blue solution. The final solution was stirred at 60°C for 1 minute before allowing the solution to cool to room temperature while continuously stirring. The particles were purified and concentrated by washing them four times with pure deionized water through a 30 kDa pressure-driven ultrafiltration system. The particles were concentrated to a peak optical extinction value of 56 $cm^{-1}$ at 720 nm (Supplementary Fig. 5c).

**Solution of Prussian blue mixed with 1% Intralipid and de-ionized water used in Fig. 3.** We used 0.47 ml Prussian blue diluted by 380 ml de-ionized water prior to adding 1% Intralipid. The measured absorption coefficient of the mixed Prussian blue and water is shown in Supplementary

Fig. 5d. This spectrum is used to predict $\mu_{eff}$ of the final solution by incorporating the measured $\mu_s$' of pure Intralipid in Ref. [3] (See Eqs. (12) and (13) in Ref. [3]). Accordingly, using a least-squares fitting to Mie theory, the measured scattering coefficient (denoted as $\mu_s$) and the anisotropy coefficient (denoted as $g$) of 10% Intralipid are [3]

$$\mu_s(\lambda) = 0.016\lambda^{-2.4} \quad (\pm 6\%)$$
$$g(\lambda) = 1.1 - 0.58\lambda \quad (\pm 5\%) \tag{S3}$$

respectively, where $\lambda$ is the wavelength between 0.4 μm and 1.1 μm and $\mu_s$ is in $mL^{-1} Lmm^{-1}$. The uncertainties of both values are also indicated [3]. Using the relations $\mu_s' \triangleq \mu_s(1-g)$ and $\mu_{eff} \triangleq \sqrt{3\mu_a\mu_s'}$ (where $\mu_a$ denotes the absorption coefficient), the predicted $\mu_{eff}$ is shown in Fig. 3c.

**Needle absorption spectrum.** Needle-weighted PA imaging requires the reference needle absorption spectrum. For this purpose, it was estimated in de-ionized water by sweeping the wavelength from 715 to 875 nm (Supplementary Fig. 5e). The needle was 21 gauge with 0.82-mm outer diameter (21G1, BD, New Jersey, NJ, USA). The estimated spectrum shown in Supplementary Fig. 5e was used as the reference spectrum in Fig. 4.

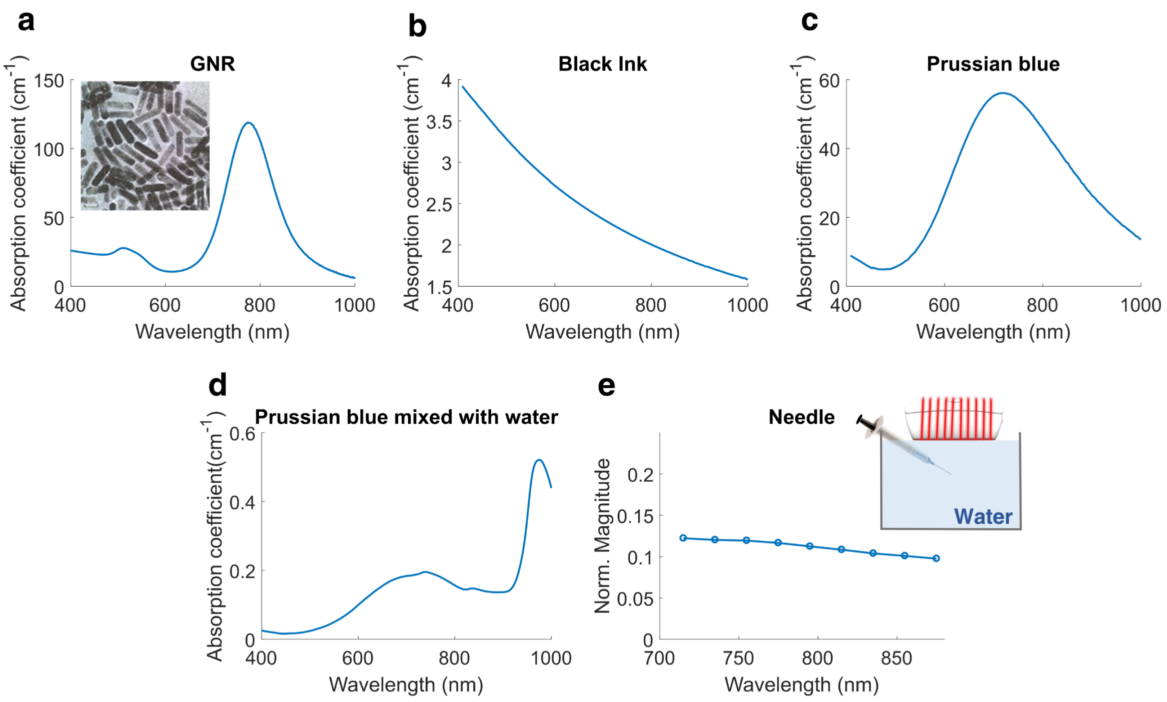


**Suppl. Fig. 5: UV-VIS and PA results for all agents used in this study**. **a,** GNR. **b,** Black ink. **c,** Custom-made Prussian blue nanoparticles. **d,** Solution used to study fluence compensation in Fig. 3 without adding Intralipid® (i.e., Prussian blue mixed with de-ionized water). **e,** Setup to measure the needle absorption spectrum in de-ionized water. The measured needle spectrum (normalized to the area under the spectrum) over the wavelength range of interest in this study.

**Supplementary Note 6**. **Light-diffusion based laser fluence estimation in the fast-sweep geometry**

For the $i$-th fiber, the reconstructed PA sub-image $S_i(\vec{r},\lambda)$ at position $\vec{r}$ and wavelength λ resulting from illumination of a short laser pulse onto an absorber inside tissue can be represented as

$$S_i(\vec{r},\lambda) = \Gamma\mu_a(\lambda)\Phi_i(\vec{r},\lambda)\ , \tag{S4}$$

where $\Gamma$ is the Grüneisen coefficient, $\mu_a(\lambda)$ is the absorber's wavelength dependent optical absorption coefficient, and $\Phi_i(\vec{r},\lambda)$ denotes the optical fluence generated by the $i$-th fiber at this position and wavelength. By coherent summation over all PA sub-images for all fibers $N_f$, the resulting PA image $P(\vec{r},\lambda)$ is expressed as

$$P(\vec{r},\lambda) = \sum_{i=1}^{N_f} S_i(\vec{r},\lambda) = \Gamma\mu_a(\lambda)\sum_{i=1}^{N_f}\Phi_i(\vec{r},\lambda)\ . \tag{S5}$$

From Eq.(S5), $P(\vec{r},\lambda)$ is simply proportional to the product of the local light absorption coefficient with the sum of laser fluences associated with integrated illumination from all 20 fibers. Hence, to accurately estimate $\mu_a(\lambda)$ from quantitative PA measurements, the fluence from each fiber, $\widehat{\Phi}_i(\vec{r},\lambda)$, must be obtained. The final fluence-compensated PA image $P_{fc}(\vec{r},\lambda)$ becomes

$$P_{fc}(\vec{r},\lambda) = \frac{P(\vec{r},\lambda)}{\sum_{i=1}^{N_f}\widehat{\Phi}_i(\vec{r},\lambda)} \cong \Gamma\mu_a(\lambda)\ . \tag{S6}$$

The fluence $\widehat{\Phi}_i(\vec{r},\lambda)$ for each wavelength and fiber illumination is estimated using the diffusion approximation (see below) to the light transport equation [4-7]. This approximation accurately models the energy distribution in a turbid medium at depths ≳ the photon transport mean free path $l_{tr} = 1/\mu_s'$ [8]. The solutions for plane, cylindrical and point sources have been presented in the literature [7-10].

The diffusion model has two independent parameters: the effective attenuation coefficient $\mu_{eff}$ and the reduced light scattering coefficient $\mu_s'$. Optimal estimates of these parameters can be obtained at a given wavelength by minimizing the mean square error between measured PA signals $\widetilde{\Phi_i'}(\vec{r},\lambda)$ from each fiber normalized to the sum of all fibers, and the normalized fluence $\Phi_i'(\vec{r},\lambda)$ predicted by the diffusion model for all $N_f$ fibers. That is

$$\underset{\mu_{eff},\mu_s'}{\operatorname{argmin}} \sum_{i=1}^{N_f}\frac{(\widetilde{\Phi_i'}(\vec{r},\lambda)-\Phi_i'(\vec{r},\lambda))^2}{N_f}\ . \tag{S7}$$

Instead of directly solving this non-linear optimization, we use brute force search to find the optimal $\mu_{eff}$ and $\mu_s'$ by confining them to a reasonable region.

The steady-state diffusion equation for an optical source $I(\vec{r},\lambda)$ describing the optical fluence distribution at depths $\gtrsim l_{tr} = 3D = 1/\mu_s'$ reduces to [4-7]

$$\mu_a(\lambda)\Phi(\vec{r},\lambda) - D(\lambda)\,\nabla^2\Phi(\vec{r},\lambda) = I(\vec{r},\lambda)\ , \tag{S8}$$

where $D(\lambda)$ is the diffusion coefficient, assumed to be spatially invariant. The exact expression of $D$ and its dependence on light absorption, scattering and anisotropy factor is still an open issue in the literature [7, 8, 11-13]. The most general expression can be found in Ref. [14]. However, in our case of $\mu_a \ll \mu_s'$ for all media used in the current work, so the diffusion coefficient does not depend on light absorption, i.e. $D = 1/3\mu_s'$ [7, 8, 14].

As shown in Supplementary Fig. 6, considering a pencil beam (i.e. $I(\vec{r},\lambda) = \delta(\vec{r})$) incident on tissue with an angle of $\theta_i$ with respect to the normal direction of the surface, the light fluence can be approximated by the illumination from two isotropic dipole sources [9, 15] and is governed by the effective light attenuation coefficient $\mu_{eff} = \sqrt{3\mu_a\mu_s'}$ and the diffusion coefficient $D$ [15-17]:

$$\Phi_i(\vec{r},\lambda) = \gamma(\lambda)\left(\frac{1}{4\pi D(\lambda)|\vec{r}-\overrightarrow{r_{p,\iota}}|}e^{-\mu_{eff}(\lambda)|\vec{r}-\overrightarrow{r_{p,\iota}}|} - \frac{1}{4\pi D(\lambda)|\vec{r}-\overrightarrow{r_{n,\iota}}|}e^{-\mu_{eff}(\lambda)|\vec{r}-\overrightarrow{r_{n,\iota}}|}\right) \quad , (S9)$$

where $\overrightarrow{r_{p,\iota}}$ and $\overrightarrow{r_{n,\iota}}$ are the position vectors of the positive and negative sources for the $i$-th fiber, respectively. Parameter $\gamma(\lambda)$ is a wavelength-dependent scaling factor.

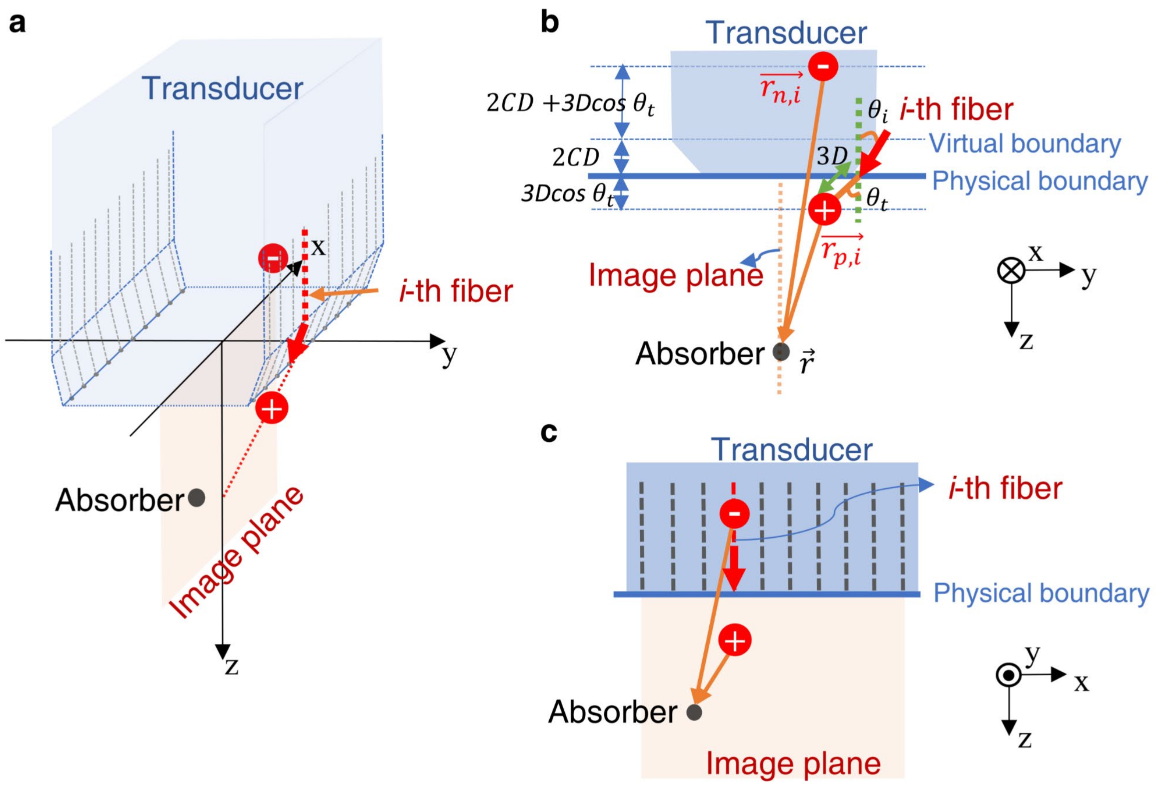


**Suppl. Fig. 6:** Geometry of light diffusion in the fast-sweep approach. The solution for $\Phi_i(\vec{r},\lambda)$ can be obtained with two positive and negative sources(indicated as positive and negative signs) [7, 9, 10, 19]. **a,** 3-D schematic view. **b,** y-z plane view. **c,** x-z plane view.

The negative source is used to satisfy the boundary condition where light propagates into the scattering medium from a non-scattering medium [16, 17]. The two sources are symmetric about the "virtual" boundary above the physical boundary by the distance $2CD$ [18]. Parameter $C$ accounts for the effects on the incident laser beam from refractive-index mismatch at the boundary, and is a function of the effective reflection coefficient $R_{eff}$ [7, 19]:

$$C \triangleq \frac{1+R_{eff}}{1-R_{eff}}\,. \tag{S10}$$

Coefficient $R_{eff}$ is zero and, therefore, $C$=1 when two media have the identical refractive index. In our application, $C = 1.04$ is determined by the refractive index mismatch between the protection glass (BK-7 optical glass) at the end of the fibers and the scattering medium.

Since both $\mu_{eff}$ and $D$ are functions of both $\mu_a$ and $\mu_s'$, three unknowns ($\gamma$, $\mu_{eff}$ and $\mu_s'$) must be estimated. Normalizing each fluence by the sum of all fiber fluences reduces the unknowns to $\mu_{eff}$ and $\mu_s'$, i.e.,

$$\Phi_i'(\vec{r},\lambda) = \frac{\Phi_i(\vec{r},\lambda)}{\sum_{i=1}^{N_f}\Phi_i(\vec{r},\lambda)}\,. \tag{S11}$$

Eq. (S11) is used for the optimization expressed in Eq. (S7).

When the diffusion coefficient $D$ is much smaller than the absorber depth, the diffusion equation can be approximated as

$$\Phi_i(\vec{r}) = -\gamma\frac{\partial}{\partial r}\frac{1}{4\pi D|\vec{r}|}e^{-\mu_{eff}|\vec{r}|}\,dr\,, \tag{S12}$$

where wavelength dependency is omitted for simplicity. We use a Cartesian coordinate system with origin at the intersection of the line between two sources and the physical boundary ($\vec{r}$ is the vector between the origin and the absorber with distance $r = \sqrt{x^2+y^2+z^2}$ ). Using the relation $dr = r^{-1}zdz$, Eq. (S12) reduces to

$$\Phi_i(\vec{r}) = \gamma\frac{zdz}{4\pi D}\frac{1}{r^3}\left(1+\mu_{eff}r\right)e^{-\mu_{eff}\cdot r}\,. \tag{S13}$$

Note that $dz = 6Dcos\theta_t + 4CD$, as indicated in Supplementary Fig. 6b. Eq. (S13) is further reduced to

$$\Phi_i(\vec{r}) = \rho\frac{z}{r^3}\left(1+\mu_{eff}r\right)e^{-\mu_{eff}\cdot r}\,, \tag{S14}$$

where $\rho = \gamma\frac{3cos\theta_t+2C}{2\pi}$. It is clear from Eq. (S14) that the light fluence is related only to the wavelength-dependent $\mu_{eff}$ independent of the diffusion coefficient. Note that the fluence is not simply a pure exponentially decaying function [16].

**Supplementary Note 7. Wavelength-dependent fluence estimation in 1%, 2%, and 4% Intralipid**

To confirm that optical fluence can be estimated using the diffusion approximation applied to PA data from the fast-sweep optical-fiber configuration, we conducted a set of experiments using human black hairs immersed in Intralipid solutions of different concentration. The Intralipid 20% IV fat emulsion (See Methods) is used as a tissue-mimicking optical scattering medium. Hair serves as a point-like optical absorber creating the PA signal used to estimate fluence variations as a function of wavelength. Three different concentrations, 1%, 2%, and 4% were prepared by diluting the original Intralipid 20% solution with de-ionized water in v/v.

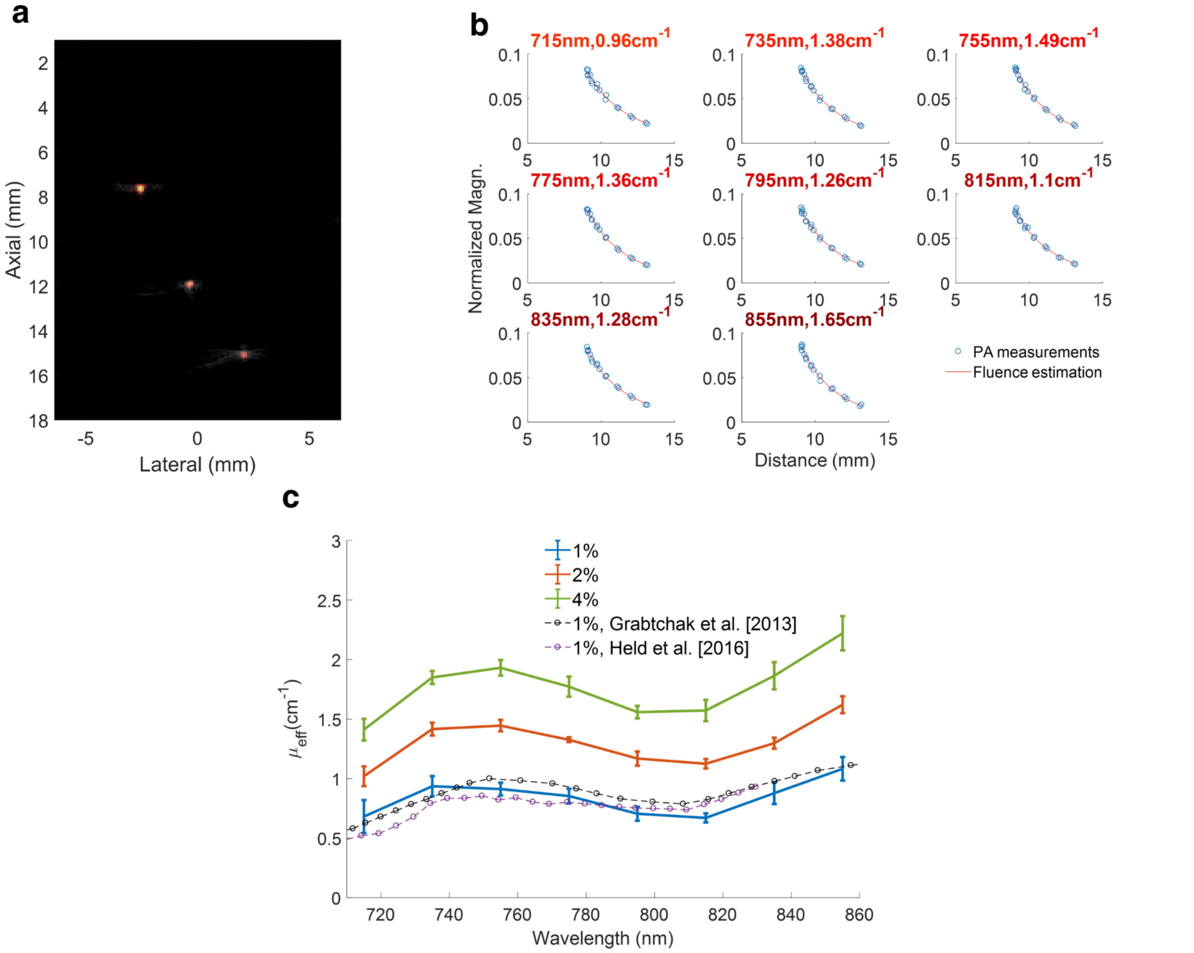


**Suppl. Fig. 7: a,** Σλ-PA image of three hairs in diluted 2% Intralipid solution. **b,** Using the hair at the shallowest depth (i.e., at 7.5 mm) for fluence estimation, the measured PA signals in the 2% solution are presented for all fibers (marked as blue circles) along with the estimated fluence (red solid lines) as a function of fiber-absorber distance (defined by the distance between a fiber and the target). For all panels, 8 different laser wavelengths ranging from 715 to 855 nm with an increment of 20 nm were evaluated. **c,** Estimated $\mu_{eff}$ (n=10) for 1%, 2%, and 4% Intralipid solutions as a function of wavelength. For comparison, two reported $\mu_{eff}$ from the literature for a 1% intralipid solution are plotted together (circled dash lines) [16, 20]. The $\mu_{eff}$ reported in [16] was 1.2% and we scaled it to 1% for comparison. The estimated $\mu_{eff}$ increases with Intralipid concentration as expected, varying approximately proportional to the square root of $\mu_s'$ when $\mu_a << \mu_s'$. Clearly, the estimated $\mu_{eff}$ in 1% Intralipid solution is consistent with results from the literature [16, 20, 21].

**<u>Supplementary Note 8</u>. E*x vivo* fluence compensation results in chicken breast**

To mimic the clinical protocol of guiding and quantifying nanoparticle injection for molecular diagnostics and therapeutics during an interventional procedure, we demonstrated fluence compensation by injecting GNRs into chicken breast. After GNR injection and needle pull out, the resulting B-mode and Σλ-PA images are shown in Supplementary Fig. 8a. Ten (10) different laser wavelengths ranging from 700 to 875 nm were used, where 700 nm was set at zero energy to estimate the noise level.

In contrast to the high scattering Intralipid medium shown in Fig. 3, chicken breast is used to mimic a relatively low scattering *in vivo* environment [18]. As shown in Supplementary Fig. 8b, the estimated $\mu_s'$ is typically under 2 cm$^{-1}$ and decreases monotonically with wavelength, generally consistent with results reported in [18]. Because $\mu_s'$ is small, the mean free path and diffusion coefficient are not small compared to the absorber depth. In contrast to Intralipid results where the laser fluence is mainly determined by $\mu_{eff}$, the laser fluence in chicken breast depends on both estimates of $\mu_s'$ and $\mu_{eff}$.

The measured spectra of GNRs were evaluated at three different lateral positions shown in the Σλ-PA image in Supplementary Fig. 8a. In each case, fluence compensation can effectively improve the GNR spectrum even though wavelength dependent fluence variations are modest in this medium compared to the Intralipid solutions.

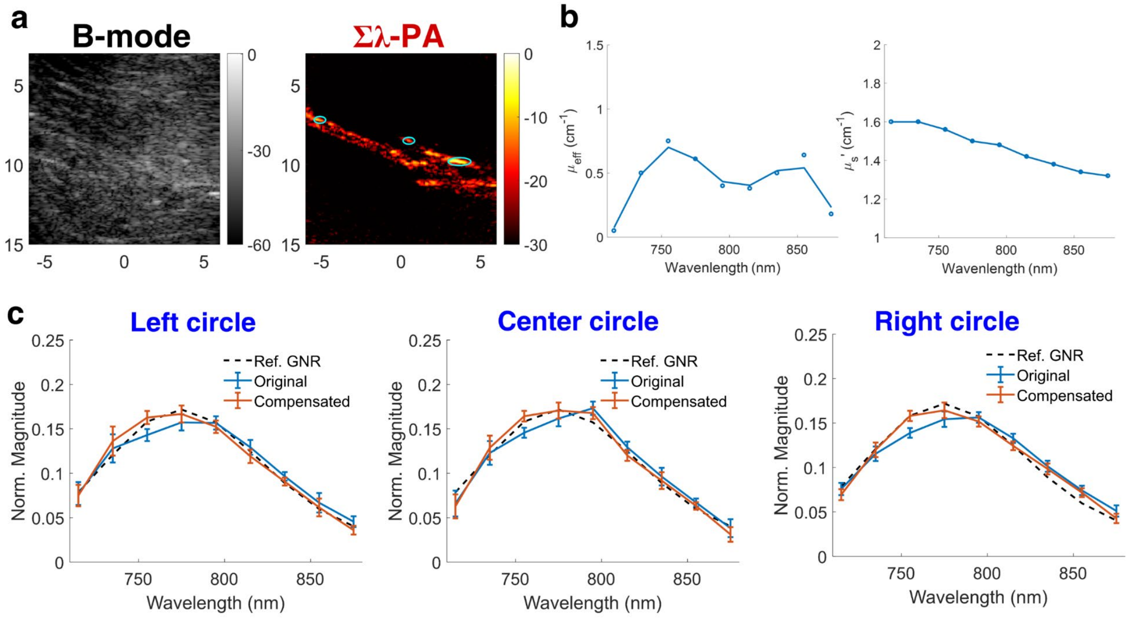


**Suppl. Fig. 8:** *Ex vivo* fluence compensation results of spectroscopic PAUS imaging for GNR injection in chicken breast. **a,** B-mode and Σλ-PA images. **b,** The estimated effective attenuation coefficient ($\mu_{eff}$, left) and reduced scattering coefficient ($\mu_s'$, right) as a function of wavelength. Both are in units of cm$^{-1}$. In the left panel, a spline-based spatial smoothing method is applied to the measured $\mu_{eff}$ (marked as circles) to produce the limited-data approximation of the function (solid line). **c,** Measured GNR spectra (normalized to the area under the spectrum) were evaluated at three different positions indicated by circles in the Σλ-PA image, statistically expressed as mean ± standard deviation. Left, middle, and right panels are the results corresponding to left, middle, and right circles. For each panel, the spectrum after fluence compensation (red solid line) is compared to the uncompensated spectrum (blue line) and the reference UV-VIS result (dashed line).

**<u>Supplementary Note 9</u>. Motion estimation using PatchMatch**

Ultrasound speckle tracking has been extensively used to estimate tissue motion [22], whether static or dynamic, and is the core signal processing method in all of US elastography [23]. It calculates all spatial components of tissue displacements from the speckle similarity between the original image frame and the frame after induced motion and deformation [24]. The most common method is a specific form of block matching, which can be realized with the multi-dimensional cross correlation function or sum of absolute differences (SAD) processing using either envelope-detected signals (phase-insensitive tracking) or complex signals (phase-sensitive tracking) [25]. Phase-sensitive tracking is preferred because tracking accuracy has been shown to be much better when phase information is included [25].

Speckle tracking produces high-resolution motion estimates because multi-dimensional correlation functions can be computed at every image pixel. When the frame rate is high, however, these computations are extensive and it is hard to produce truly real-time estimates. Recently, we introduced an algorithm based on a randomized search called PatchMatch [26] to speed up processing for near real-time and truly real-time implementation [14]. We showed that PatchMatch performance is comparable to that of an extensive search while the computation load is significantly reduced by at least 10 times [27]. In addition, PatchMatch implicitly imposes smoothness on displacement estimates in adjacent pixels, which is robust against speckle decorrelation where speckle similarity is corrupted by tissue deformation, low SNR, and other physical effects [27].

As shown here, there are three major steps in PatchMatch: initialization, propagation, and random search [26]. Take a kernel (indicated as a blue cube) in the reference volumetric image as an example. The 3-D case is presented here as the most general form of speckle tracking, but in the present study only 2-D tracking was performed on B-Mode image loops. This kernel and its spatially closest neighbors (indicated with different colors, Supplementary Fig. 9a) are assigned to the uniformly-distributed random positions in the target volumetric image (Supplementary Fig. 9b). In practice, such random assignment is confined to a pre-defined search region according to *a priori* knowledge of maximum tissue motion. The similarity between the individual pair in reference and target images is measured using the NCC (defined in Eq. (S.1)). By comparing the NCC values of all neighbors, the kernel propagates to the neighbor with maximum NCC value (Supplementary Fig. 9c). To avoid falling into a local maximum, additional search is performed by randomly selecting other positions (Supplementary Fig. 9d), confined to a pre-defined search region. In this study, six random positions are empirically determined. Last, the final position in the target image is determined by calculating the maximum NCC among all seven positions (Supplementary Fig. 9e). By iterating between propagation and random search, the final positions for all kernels converges efficiently (Supplementary Fig. 9f). In practice, iteration is performed in windshield-wiping order, i.e., from left/top to right/down and then scanned backward. Thus, two iterations complete one scan cycle. Accordingly, 2-D displacements between two US images are estimated. We have shown that 4 iterations are enough to obtain reliable estimates [27].

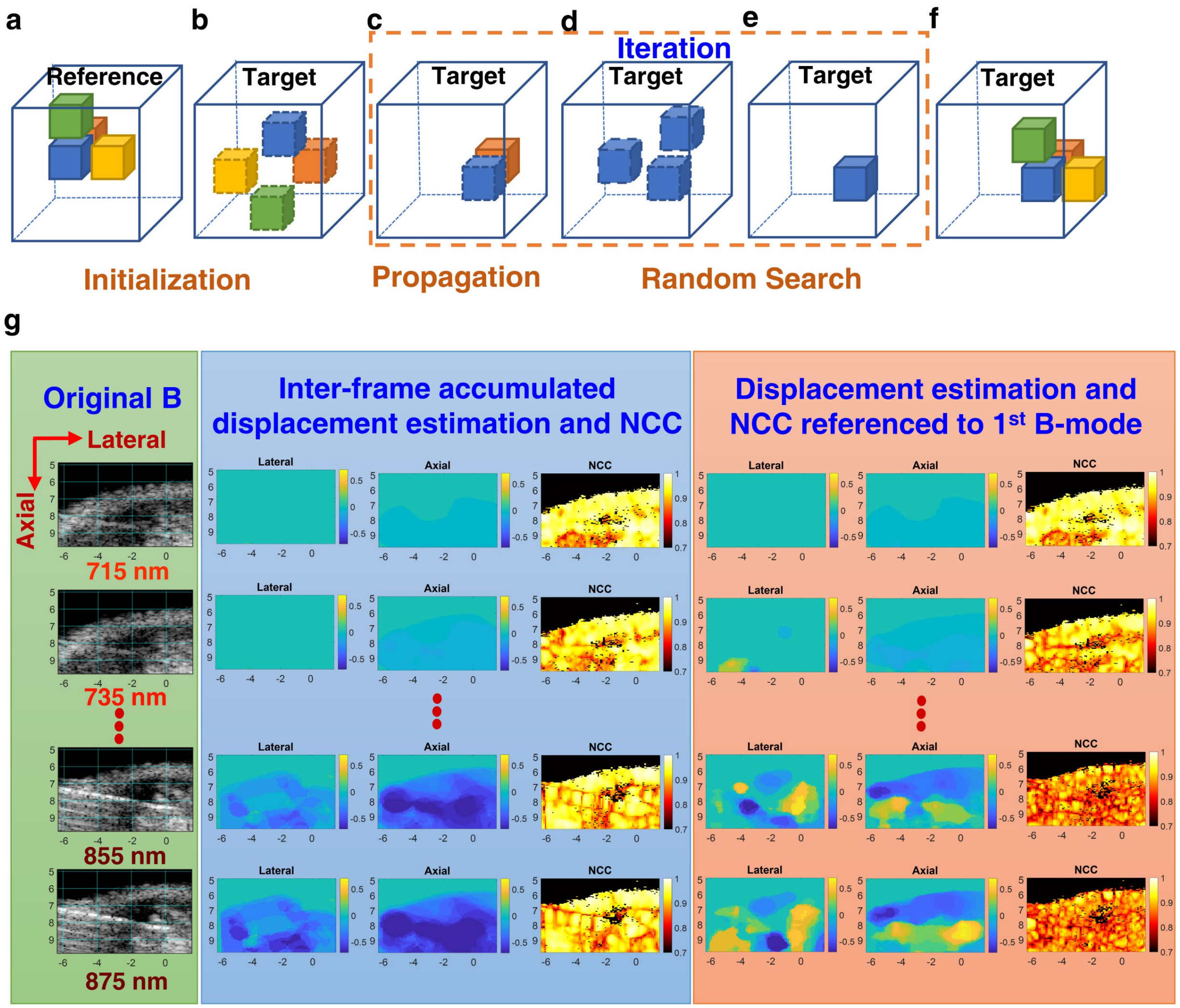


**Suppl. Fig. 9.** Schematic flow chart of PatchMatch for multi-dimensional displacement estimation (3-D estimation is shown here as the most general case – only 2-D tracking was performed in this study) [26, 27]. Taking a kernel (indicated as a blue cube) in the reference image as an example, **a,** six closest neighbors surrounding this kernel (only three are displayed) are indicated with different colors. **b,** The positions of the kernel and its neighbors in the target image are randomly assigned. The similarity between individual pairs in reference and target images (i.e., identical color cubes) is measured. **c,** The kernel propagates to the neighborhood having the maximum similarity. **d,** Other random positions are chosen to avoid a local maximum. **e,** Determine the final position by comparing the similarity of all candidates in (d). **f,** Iteration is performed between (c)-(e) to find the positions of all kernels. It has been shown that 4 iterations are sufficient in ultrasound motion estimation. Accordingly, displacements between two images are estimated. **g,** *in vivo* PatchMatch-based speckle tracking results of needle guidance in a mouse. Uncompensated B-mode images associated with the laser emission from 715 to 875 nm are shown in the left column. Motion estimation (i.e., lateral and axial displacement components) between two successive B-mode images (i.e., interframe or inter-wavelength estimation, middle column) is compared with that using the first B-mode (i.e., at 700 nm wavelength) as a reference (right column). Note that accumulated displacement is required for interframe estimation. As an index of tracking reliability, the NCC map for each tracking result is also presented.

Supplementary Fig. 9g shows *in vivo* results using PatchMatch-based speckle tracking on the 10 B-mode images corresponding to 10 laser wavelengths ranging from 700 to 875 nm. As mentioned earlier, we disabled 700 nm laser emission to estimate the noise level but still acquired B-mode images at this wavelength for tracking purposes. Original B-mode images corresponding to 715 nm to 875 nm are shown in the left column (same as in Fig. 4). When comparing different B-mode images, it is clear that speckle decorrelation is significant because of the appearance of a needle. Interframe (or equivalently, inter-wavelength) displacement estimation results (i.e., estimation between two successive images, middle column) are compared to the results using the first image as a reference (right column). Note that interframe estimation requires accumulation of all previous paired displacement estimates. Lateral and axial displacement components are shown in units of mm. Positive values for lateral/axial components represent the right/down direction, respectively. The NCC map for each tracking result is also shown as a reliability index. In the first two images, the two estimation approaches are similar because speckle decorrelation is small. However, for 855 and 875 nm, last-to-first frame displacement estimation exhibits larger errors as both lateral and axial components have diverse motion directions. This is because speckle decorrelation is significant, as evidence by the much smaller NCC values for this case. In contrast, interframe displacement estimation produces higher NCC values and uniform 2-D motion estimates under high speckle decorrelation, which is more robust than last-to-first frame estimated displacements.

The results in Supplementary Fig. 9g show that inter-wavelength (i.e., inter-frame) displacement estimation can maintain high speckle correlation with high frame rates (i.e., 50 Hz), enabling robust speckle tracking. However, if low frame rate images are used for motion compensation, as shown in Supplementary Fig. 9g where the equivalent frame rate from 1$^{st}$ to last wavelength is around 5 Hz, speckle decorrelation worsens displacement estimation. Therefore, motion compensation is only realistic with high frame rate imaging.

When tissue motion changes rapidly with time (e.g., cardiac applications), the flexibility of our imaging pulse sequence enables interleaved PAUS images where the PA frame rate is 50 Hz but the US frame rate can be much higher by incorporating high speed US imaging techniques such as parallel/multibeam and compounded plane waves with a small number of angles. Increasing US frame rate to track fast motions enables more robust motion estimation while still preserving the overall 50 Hz interleaved PAUS imaging rate.

### Supplementary movie captions

**Supplementary Movie 1**: Real-time *ex vivo* spectroscopic PAUS imaging for needle guidance with nanoparticle injection at a 50-Hz frame rate showing three stages: needle insertion into chicken breast, GNR injection, and needle pullout. For each stage, PA images are acquired at a single wavelength (775 nm), followed by a 10 wavelength sweep ranging from 700 to 875 nm. The laser sequence (right) is shown synchronized with the real-time image (left).

**Supplementary Movie 2**: Real-time *in vivo* mouse spectroscopic PAUS imaging for needle guidance with nanoparticle injection at a 50-Hz frame rate. PA images are acquired at a single wavelength (775 nm), followed by a 10 wavelength sweep ranging from 700 to 875 nm. The laser sequence (right) is shown synchronized with the real-time image (left).

### Supplementary References


1. Zhu, W., Liu, K, Sun, X, Wang, X., Li, Y., Cheng, L., Liu, Z. Mn2+-doped prussian blue nanocubes for bimodal imaging and photothermal therapy with enhanced performance. *ACS Appl. Mater. Interfaces* **7**(21), 11575-11582 (2015).
2. Ren, W., Qin, M., Zhu, Z., Yan, M., Li, Q., Zhang, L., Liu D., Mai, L. Activation of Sodium Storage Sites in Prussian Blue Analogues via Surface Etching. *Nano Lett.* **17**(8), 4713-4718 (2017).
3. Van Staveren, H. J., Moes, C. J. M., van Marie, J., Prahl, S. A., van Gemert, M. J. C. Light scattering in lntralipid-10% in the wavelength range of 400–1100 nm. *Appl. Opt.* **30**(31), 4507-4514 (1991).
4. Van de Hulst, H.C. *Light Scattering by Small Particles* (Dover Publications, New York, 1981).
5. Furutsu, K., Yamada, Y. Diffusion approximation for a dissipative random medium and the applications. *Phys. Rev. E* **50**(5), 3634 -3640 (1994).
6. Ishimaru, A. *Wave propagation and scattering in random media* (Academic Press, New York, 1977).
7. Karabutov, A. A., Pelivanov, I. M., Podymova, N. B., Skipetrov, S. E., Determination of the optical characteristics of turbid media by the laser optoacoustic method. *Quantum Electronics* **29**, 215-220 (1999).
8. Grashin, P. S., Karabutov, A. A., Oraevsky, A. A., Pelivanov, I.M., Podymova, N. B., Savateeva, E. V., Solomatin, V. S. Distribution of the laser radiation intensity in turbid media: Monte-Carlo simulations, theoretical analysis and the results of optoacoustic measurements. *Quantum Electronics* **32**, 868-874 (2002).
9. Jacques, S. L. Light distributions from point, line and plane sources for photochemical reactions and fluorescence in turbid tissues. *Photochem. Photobiol.* **67**(1), 23-32 (1998).
10. Jia, M., Chen, X., Zhao, H., Cui, S., Liu, M., Liu, L., Gao, F. Virtual-source diffusion approximation for enhanced near-field modeling of photon-migration in low-albedo medium. *Opt. Express* **23**(2), 1337-1352 (2015).
11. Corngold, N., Aronson, R. Photon diffusion coefficient in an absorbing medium. *J. Opt. Soc. Am.* **16**(5), 1066-1071 (1999).
12. Pierrat, R., Greffet, J.-J., Carminati, R. Photon diffusion coefficient in scattering and absorbing media. *J. Opt. Soc. Am.* **23**(5), 1106-1110 (2006).
13. Ripoll, J., Yessayan, D., Zacharakis,G., Ntziachristos, V. Experimental determination of photon propagation in highly absorbing and scattering media. *J. Opt. Soc. Am.* **22**(3), (2005).
14. Glasston, S., Edlund, M. C. *The elements of nuclear reactor theory* (D. Van Nostrand Co., Inc., New York, 1952).

15. Farrell, T.J., M.S. Patterson, and B. Wilson, A. diffusion theory model of spatially resolved, steady-state diffuse reflectance for the noninvasive determination of tissue optical properties in vivo. *Med. Phys.* **19**(4), 879-888 (1992).

16. Held, K. G., Jaeger, M., Rička, J., Frenz, M., Akarçay, H. G. Multiple irradiation sensing of the optical effective attenuation coefficient for spectral correction in handheld OA imaging. *Photoacoustics* **4**(2), 70-80 (2016).

17. Ranasinghesagara, J. C., Zemp, R. J. Combined photoacoustic and oblique-incidence diffuse reflectance system for quantitative photoacoustic imaging in turbid media. *J. Biomed. Opt.* **15**(4), 046016 (2010).

18. Marquez, G., Wang, L. V., Lin, S. P., Schwartz, J. A., Thomsen, S. L. Anisotropy in the absorption and scattering spectra of chicken breast tissue. *App. Optics*, **37**(4), 798-804 (1998).

19. Wang, L.V., Wu, H.-I., *Biomedical optics: principles and imaging*. (John Wiley & Sons, 2012).

20. Grabtchak, S., K.B. Callaghan, and W.M. Whelan, Tagging photons with gold nanoparticles as localized absorbers in optical measurements in turbid media. *Biomed. Opt. Express* **4**(12) 2989-3006 (2013).

21. Flock, S. T., Jacques, S. L., Wilson, B. C., Star, W. M., van Gemert, M. J. Optical properties of Intralipid: a phantom medium for light propagation studies. *Lasers Surg. Med.* **12**(5), 510-519 (1992).

22. Kaluzynski, K., Chen, X., Emelianov, S.Y., Skovoroda, A. R., O'Donnell, M. Strain rate imaging using two-dimensional speckle tracking. *IEEE Trans. Ultrason. Ferroelectr. Freq. Control* **48**(4), 1111-1123 (2001).

23. D'Hooge, J., Konofagou, E., Jamal, F., Heimdal, A., Barrios, L., Bijnens, B., Thoen, J., Van de Werf, F., Sutherland, G., Suetens, P. Two-dimensional ultrasonic strain rate measurement of the human heart in vivo. *IEEE Trans. Ultrason. Ferroelectr. Freq. Control* **49**(2), 281-286 (2002).

24. Lubinski, M.A., Emelianov, S.Y., O'Donnell, M. Speckle tracking methods for ultrasonic elasticity imaging using short-time correlation. *IEEE Trans. Ultrason. Ferroelectr. Freq. Control* **46**(1), 82-96 (1999).

25. Viola, F., Walker, W.F. A comparison of the performance of time-delay estimators in medical ultrasound. *IEEE Trans. Ultrason. Ferroelectr. Freq. Control* **50**(4), 392-401 (2003).

26. Barnes, C., Shechtman, E., Finkelstein, A., Goldman, D. B. PatchMAtch: a randomized correspondence algorithm for structural image editing. *ACM Transactions on Graphics* (Proc. SIGGRAPH) **28**(3), (2009).

27. Jeng, G.S., Zontak, M., Parajuli, N., Lu, A., Ta, K., Sinusas, A. J., Duncan, J. S., O'Donnell, M. Efficient two-pass 3-D speckle tracking for ultrasound imaging. *IEEE Access* **6**, 17415-17428 (2018).
